# Integrating inflammatory biomarker analysis and artificial intelligence-enabled image-based profiling to identify drug targets for intestinal fibrosis

**DOI:** 10.1101/2022.06.08.495258

**Authors:** Shan Yu, Alexandr A. Kalinin, Maria D. Paraskevopoulou, Marco Maruggi, Jie Cheng, Jie Tang, Ilknur Icke, Yi Luo, Qun Wei, Dan Scheibe, Joel Hunter, Shantanu Singh, Deborah Nguyen, Anne E. Carpenter, Shane R. Horman

**Author notes:** Corresponding Author: Address: Takeda Development Center Americas, Inc., 9625 Towne Center Dr, San Diego, CA, 92121. Shan Yu, Shane R. Horman.

## Abstract

Intestinal fibrosis is a common complication of several enteropathies with inflammatory bowel disease being the major cause. The progression of intestinal fibrosis may lead to intestinal stenosis and obstruction. Even with an increased understanding of tissue fibrogenesis, there are no approved treatments for intestinal fibrosis. Historically, drug discovery for diseases like intestinal fibrosis has been impeded by a lack of screenable cellular phenotypes. Here we applied Cell Painting, a scalable image-based morphology assay, augmented with machine learning algorithms to identify small molecules that were able to morphologically reverse the activated fibrotic phenotype of intestinal myofibroblasts under pro-fibrotic TNFα stimulus. In combination with measuring CXCL10, a common pro-inflammatory cytokine in intestinal fibrosis, we carried out a high-throughput small molecule chemogenomics screen of approximately 5000 compounds with known targets or mechanisms, which have achieved clinical stage or approval by the FDA. Through the use of two divergent analytical methods, we identified several compounds and target classes that are potentially able to treat intestinal fibrosis. The phenotypic screening platform described here represents significant improvements in identifying a wide range of drug targets over conventional methods by integrating morphological analyses and artificial intelligence using pathologically-relevant cells and disease-relevant stimuli.

## Introduction

Intestinal fibrosis is a pathophysiological mechanism of intestinal tissue repair that leads to the deposition of desmoplastic connective tissue after injury. This process can be triggered by noxious agents, including infections, autoimmune reactions, physical, chemical, and mechanical injuries. Under normal physiological conditions, intestinal immune components can help to clear foreign pathogens and facilitate tissue repair through canonical wound healing processes. However, fibrogenesis may occur when the immune response is uncontrolled and persistent, or when injuries repeat, resulting in chronic damage^*2, 3*^. Intestinal fibrosis is one of the most common complications of patients who suffer from inflammatory bowel disease (IBD), occurring in approximately 5% of ulcerative colitis (UC) patients and more than 30% of Crohn’s disease patients. The prevalence for IBD increased from 0.5% in 2010 to 0.75% in 2022 in western countries and is projected to reach 1% in 2030^*4, 5*^. Fibrostenotic complications, including stricture formation and subsequent intestinal obstruction, significantly increase morbidity and hospitalization, surgical intervention, and health care costs^*2*^. Despite advances in the development of therapeutics for treating IBD, including small molecular weight immunomodulators (prednisone, 5-aminosalycylic acid, Tofacitanib, Ozanimod), DNA/RNA replication inhibitors (azathioprine, methotrexate, 6-mercaptopurine), and large molecular weight anti-inflammatory biologics (anti-TNFα, anti-integrins and anti-IL-12/IL-23), the high incidence of intestinal strictures and requirement for surgical interventions remain^*6*^. The lack of effective drug therapies for fibrostenotic IBD represents an increasing and significant unmet medical need.

At a molecular basis, intestinal fibrosis in IBD is a dynamic and multifactorial process. It is a consequence of local chronic inflammation and subsequent activation of fibroblasts. Mucosal inflammation occurs when the mucosal integrity is compromised resulting in the influx of micro-organisms from the gut lumen. Myeloid cells, such as macrophages and dendritic cells, recognize these pathogen-associated molecular patterns via Toll-like and NOD-like pattern recognition receptors and propagate the immune signaling by recruiting other immune cells to clear the offending pathogens by releasing cytokines and chemokines, such as TNFα, IL-1β, IL-36 and OSM^*7*^. Tissue repair and wound healing occurs in the resolution of the inflammation process after initial inflammatory responses. However, in the context of chronic inflammation, cytokines and chemokines drive differentiation and activation of fibroblasts and their subsequent production of ECM proteins. When the balance between production and enzymatic degradation of ECM proteins is lost, intestinal fibrosis occurs^*6*^.

Due to the failure rate of translational efficacy for many clinical candidates for IBD^*8*^, there is an increased interest in the exploratory phase of drug discovery, to utilize disease-relevant phenotypic screening to provide more confidence to identify drug targets or small molecules^*9-11*^. However, lead molecules derived from phenotypic screening campaigns are historically difficult to follow up due to intrinsic complexities of generating useful structure-activity relationships, and lack of structure-based drug design input, coupled to the difficulties in predicting and successfully navigating mechanism-associated toxicities. Chemogenomic screening utilizes a library of selective small molecules with annotated targets. The benefit of phenotypically profiling compounds with known targets and mechanisms is to assist generation of mechanistic hypotheses that can initiate ensuing target validation studies. Furthermore, hit molecules identified from such screens can also suggest that their targets are amenable to functional pharmacological modulation, thus providing evidence of the druggability of the targets^*11*^.

Due to practicality and affordability, drug discovery campaigns typically employ one or a few readily interpretable biomarkers, such as secretory or intracellular markers or gene-of-interest-driven reporters, that reflect known biology. Recently, significant interest has arisen in the drug discovery industry to capture high-dimensional cellular morphological changes to stimuli and drug treatments by using image-based profiling with automated microscopy^*12*^. This unbiased, inexpensive and scalable image-based method, most often using the Cell Painting assay, combines multiple organelle stains in a robust assay yielding single-cell profiles composed of thousands of features^*12*^. Combined with machine learning and data mining, Cell Painting offers the potential to accelerate therapeutic discovery by identifying drug-induced cellular phenotypes, elucidating modes of action, and characterizing drug toxicities^*12*^.

In this study, we describe a chemogenomic library screen in human intestinal fibroblasts using both disease-relevant biomarker and Cell Painting readouts to interrogate targeted small molecules that can alleviate the fibrotic phenotype. We identified clinically relevant hits from both assay readouts, though the mechanisms-of-action of hits from each assay represent distinct fibrotic biology. We identified inflammatory response regulators with the biomarker assay, and tissue plasticity, remodeling, fibrosis and angiogenesis signaling modulators with the Cell Painting assay. The hits were further confirmed and validated in colonic fibroblasts treated with other pro-fibrotic stimuli. With this integrated approach using both high throughput biomarker analysis and artificial intelligence-enabled morphological profiling, we were able to discover a wide spectrum of physiologically and clinically relevant small molecules and targets for intestinal fibrosis.

## Results

### Development of an in vitro cellular disease model that mimics human intestinal fibrosis pathogenic cell population

The CCD-18co human colon fibroblast cell line was identified as a physiologically relevant model for human intestinal fibroblasts^*13*^. In order to identify culture conditions that yielded the most clinically relevant response to disease-associated stimuli, we performed single cell transcriptomic analysis of CCD-18co cells that were treated with various pro-fibrotic stimuli, including TNFα, IL-1β, TGFβ, TL1a, OSM and IL-36, for 16 hours. We combined data from each treatment in an integrated UMAP (Figure 1A) and compared their single-cell RNA sequencing profiles side-by-side (Figure 1B). We identified seven distinct clusters of cells in total, of which several common clusters were shared among all treatments, as well as unique clusters corresponding to particular treatment groups (Figure 1B).

**Figure 1.**
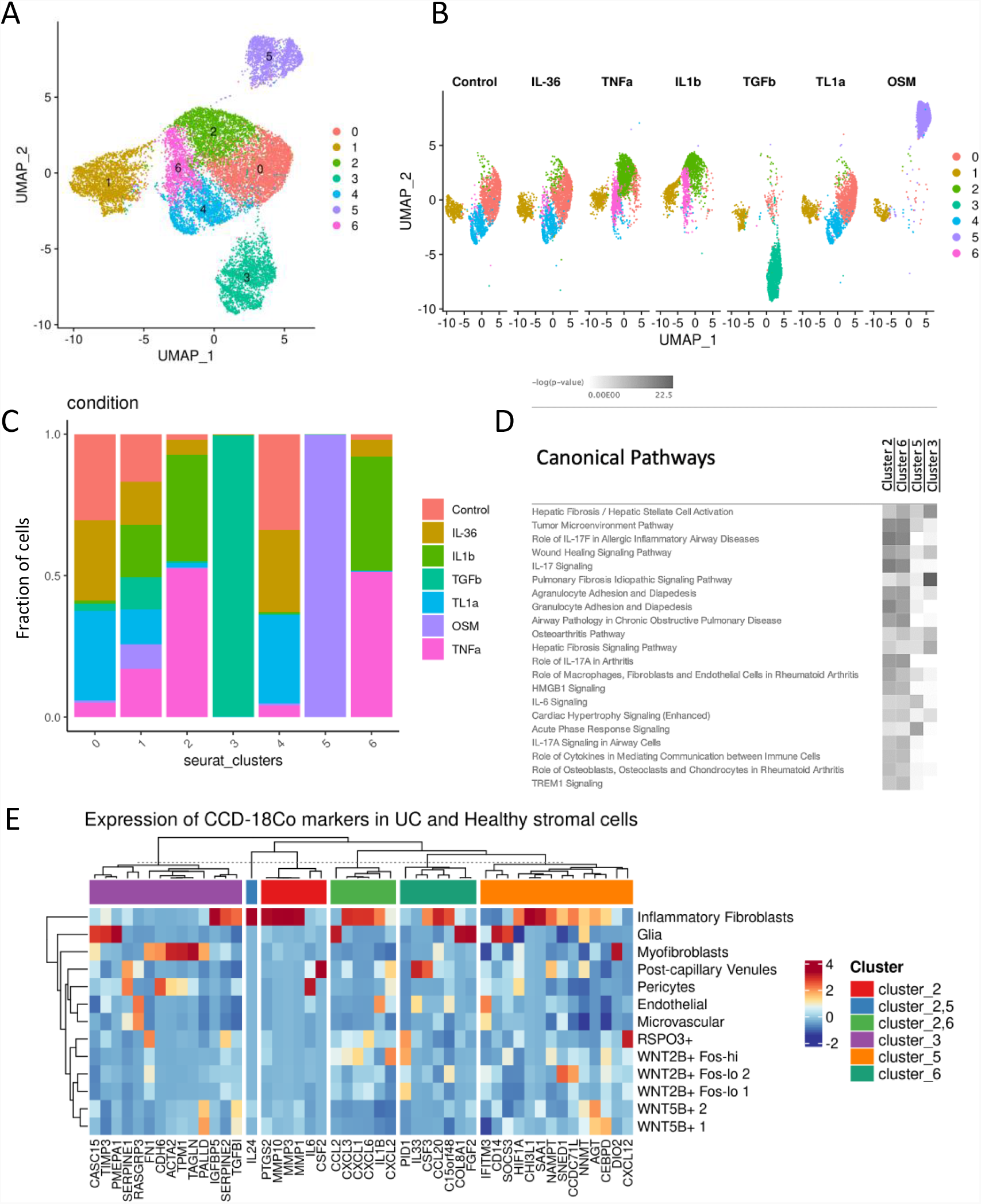
Bioinformatic analysis of transcriptome profile of CCD-18co cells and comparison with human colon biopsies. (A) and (B) UMAP embedding of 16,750 single-cell RNA sequencing (scRNA-Seq) profiles from CCD-18co fibroblast cell cultures with different stimuli, including TNFα, IL-1β, TGFβ, TL1a, OSM and IL-36, for 16 hours. Seven identified single-cell clusters are indicated by colors. (C) Stacked bar graph showed cluster cell composition, with cluster 3 corresponding to cells activated by TGFβ, cluster 5 corresponding to cells stimulated by OSM, while clusters 2 and 6 captured cellular responses upon TNFα and IL-1β treatments. The remaining clusters were not overrepresented in any of the conditions and were considered baseline state. (D) Ingenuity Pathway Analysis (IPA)Canonical Pathways associated with the upregulated genes in clusters 2 and 6 (TNFα and IL-1β stimuli), cluster 5 (OSM) and cluster 3 (TGFβ). Cluster 1, not shown in Figure 1D, exhibited high expression of cell cycle phase genes. Grayscale represents p-score = − log10 (p-value). (E) Top 15 markers from CCD-18co fibroblast cell clusters 2, 3, 5, and 6 were analyzed in human colon fibroblasts from UC and healthy patients, retrieved from the published stromal single cell atlas ^*1*^. Highly expressed genes in CCD-18co cluster 2, 5, 6 (TNFα, OSM, IL-1β treatments) were enriched in Inflammatory Fibroblasts, and highly expressed genes in CCD-18co cluster 3 (TGFβ cellular treatment) were elevated in Myofibroblasts from colonic biopsies.

Within these clusters we performed functional characterization by mapping the enriched canonical pathways and upstream regulators. Clusters 2 and 6 were predominant in TNFα and IL-1β treatment groups (Figure 1C). Genes upregulated in these clusters represented IL-17 signaling, wound healing, TREM1 signaling, cytokine mediated fibroblast crosstalk, leukocyte migration and tumor microenvironment pathways, as well as genes involved in mediating inflammatory pathways associated with cancer (Figure 1D). Cluster 3 and cluster 5 were mainly found in TGFβ and OSM treatments, respectively (Figure 1C). Genes upregulated in cluster 5 represented IL-6 signaling and acute phase response signaling, while genes upregulated in cluster 3 represented tissue fibrosis activities (Figure 1D). IL-36 and TL1A treatment profiles were similar to control, suggesting neither stimulus exerted significant effect on the cells (Figure 1B). Upstream regulator detection analysis corroborated that the clusters 2,6 are modulated by TNFα and IL-1β, while cluster 3 from TGFβ and cluster 5 from OSM.

To identify which CCD-18co population exhibited the most disease-mimetic gene expression profile, we mapped activated CCD-18co clusters (cluster 2, 6, 3 and 5) to cell populations from primary human colon stromal biopsies from healthy and ulcerative colitis patients^*1*^ (Figure 1E). We found that cluster 2 and 6, most prevalent in TNFα and IL-1β treatments, and cluster 4, unique to OSM treatment, had signatures that closely overlapped with those of inflammation-associated fibroblasts (IAFs) in diseased human colon biopsies. Cluster 3, specific to TGFβ treatment, corresponded to both IAFs and myofibroblasts in human colon biopsies. Because IAFs are the immunological hub of multiple signaling pathways that play important roles during the onset of intestinal inflammation and fibrosis^*14*^, and IAFs are associated with anti-TNFα drug resistance in inflammatory bowel disease patients^*1*^, we sought to address this key unmet medical need for intestinal fibrosis and perform the primary screen with TNFα as stimulus, as it was found to induce an IAF phenotype.

To quantify the effects of TNFα signaling on morphological fibrosis in CCD-18co cells, we knocked out *TNFRSF1A* and *TNFRSF1B* genes, which encode TNFR1 and TNFR2, respectively, individually or together using CRISPR/Cas9 gene editing (S. Figure 1A), then evaluated the response of the cells to TNFα. Upon activation of NF-kB by TNFα signaling, p65, a subunit of NF-kB also known as RELA, was observed to translocate from the cytoplasm to the nucleus (S. Figure 1B). However, cells transfected with individual or pooled TNFRSF1A guide RNAs (gRNAs) showed that p65 remained, at least partially, in the cytoplasm (S. Figure 1C), indicating reduced NF-kB signaling. Further, CCD-18co cells transfected with individual or pooled TNFRSF1A gRNAs showed diminished CXCL10 expression compared to control cells (S. Figure 1D). The effect of dual TNFRSF1A and TNFRSF1B knockout was similar to TNFRSF1A knockout alone indicating TNFα signaling was mediated, at least partially, through TNFR1 instead of TNFR2 in CCD-18co cells.

In high-throughput screening, it is important to use clinically proximal readouts whenever possible to ensure the observed phenotype is a robust surrogate for disease pathology. To that end, we assessed protein and mRNA expression levels of a panel of inflammation-related biomarkers in CCD-18co cells that were treated with disease-relevant pro-fibrotic stimuli. We identified CXCL10 as a significantly upregulated biomarker at both protein and mRNA levels by multiple stimuli, including TNFα, IL-1β and IL-36 (S. Figure 2). Because modulation of CXCL10 and its receptor CXCR3 has been reported to be associated with inflammatory signaling-driven fibrogenesis^*15, 16*^, we chose it as a readout for efficacy in the ensuing screen. Though we did profile more conventional biomarkers of fibrosis, including ACTA2 and COL1A1, neither of them was induced by pro-fibrotic stimuli at either protein or mRNA level to yield an acceptable assay window for a high throughput screen (S. Figure 3).

**Figure 2.**
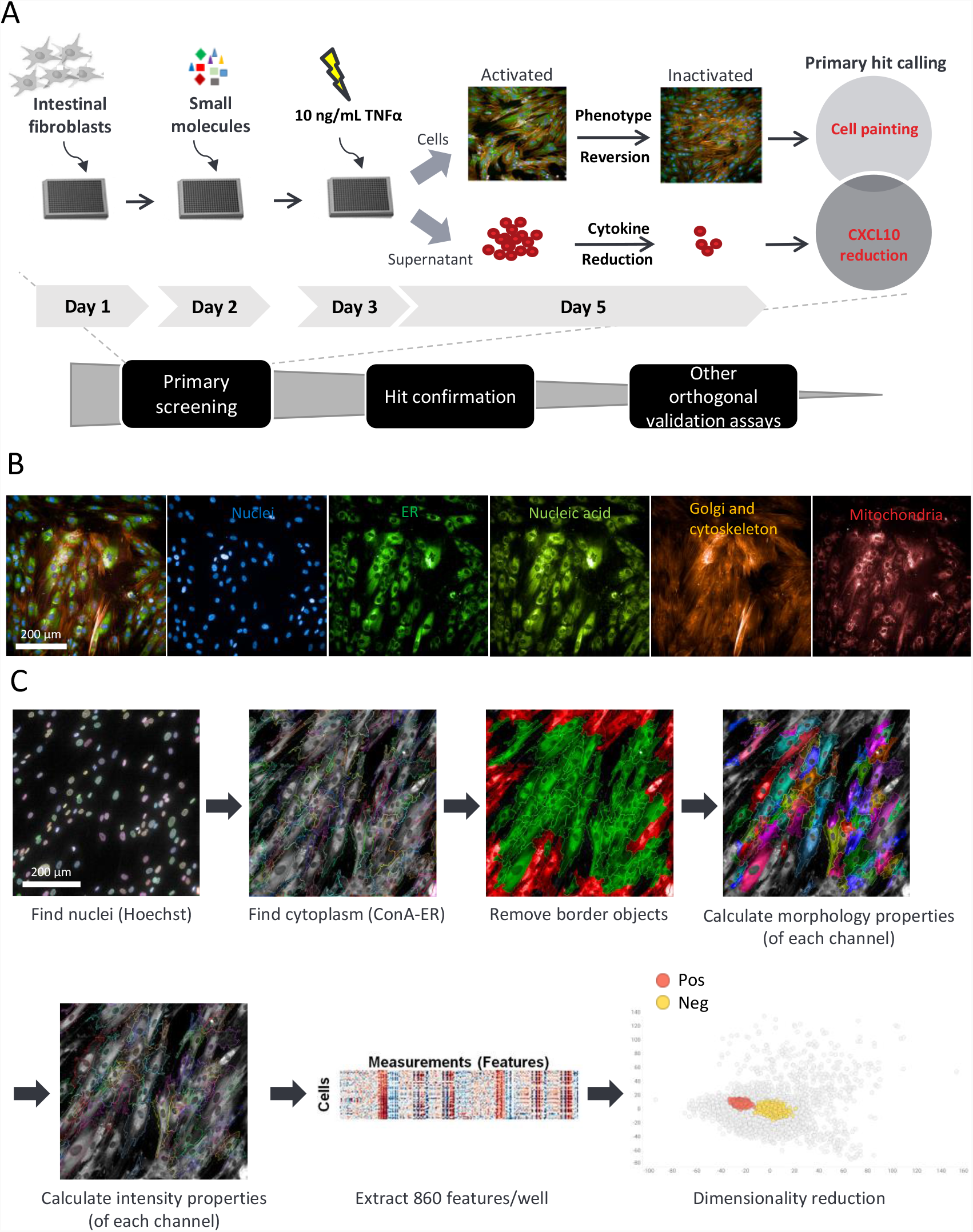
Chemogenomic library screen workflow. (A) The screen was conducted through a process including primary screening, hit confirmation and other orthogonal validation assays. For the primary screen, colonic fibroblasts CCD-18co cells were plated on Day 1, followed by small molecule transfer on Day 2, and 10 ng/mL TNFα stimulation on Day 3. The supernatant samples were collected for the CXCL10 reduction assay and cells were stained with the Cell Painting dyes for the high content imaging assay. Hits from both assays were called and analyzed individually and collectively (B) CCD-18co cells that were stained with Cell Painting dyes including Hoechst 33342 (nuclei), Concanavalin A-Alexa 488 (ER), SYTO™ 14 (nucleic acid), WGA-Alexa 555 (Golgi), phalloidin-Alexa 568 (cytoskeleton) and MitoTracker Deep Red (mitochondria), and imaged with Operetta CLS. The image on the far left was merged image of all channels. (C) Workflow of cellular compartment segmentation of high content images using Perkin Elmer Harmony software. Nuclei were identified by Hoechst 33342 stain. Cytoplasm was then identified by Concanavalin A-Alexa 488 stain. The border objects were excluded from analysis. Different morphology and intensity properties of each channel were calculated and 860 features were extracted at the well-level. The profiling dataset was then analyzed with a dimensionality reduction method, such as PCA.

**Figure 3.**
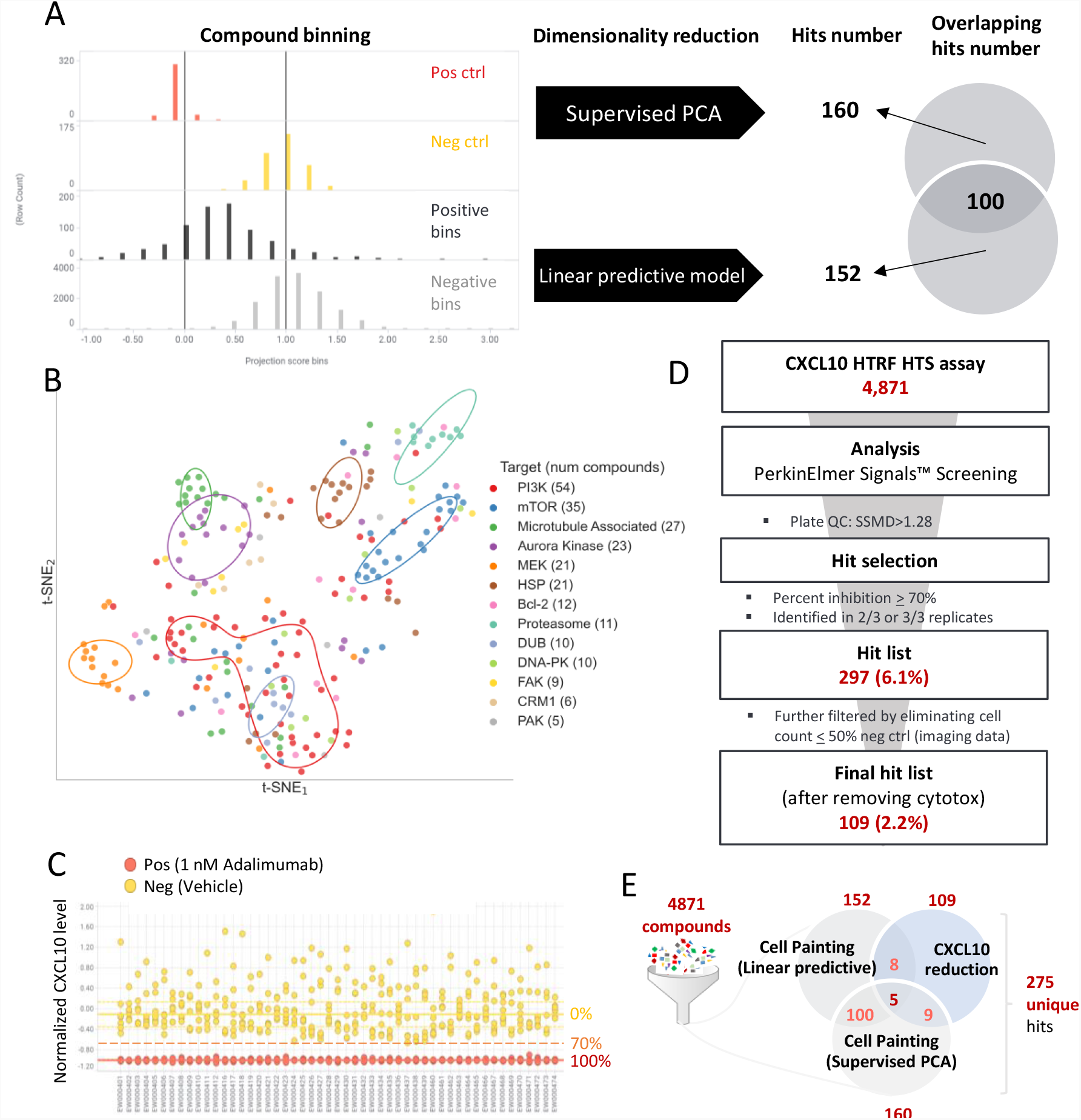
Primary screen hit picking strategies for the CXCL10 reduction assay and Cell Painting assay. (A) The Cell Painting dataset was analyzed with both supervised PCA and linear predictive model methods. Projection scores of Cell Painting controls and samples help to determine the similarities between compounds and controls. Compounds in positive bins in the range between Average(projection score) ± 3 x S.D. were picked as hits. There were 160 hits from supervised PCA and 152 hits from linear predictive model analysis with 100 overlapping hits. (B) t-SNE plot shows the phenotypic space of top compound target categories that are farthest from the negative controls. (C) Pos and neg ctrl data points of CXCL10 HTRF assay. X-axis shows the plate barcode, y-axis shows the normalized CXCL10 level. Solid yellow line shows 0% inhibition representing the median of the neg ctrl (vehicle), and solid red line shows 100% inhibition representing the median of pos ctrl (1 nM adalimumab). Dotted orange line shows 70% cut off for hit picking. (D) The CXCL10 HTRF assay screened 4,871 compounds in total. The plates passed through QC criteria of SSMD>1.28. Hits were selected if 2 or 3 out of the triplicates exhibit percent inhibition of CXCL10 ≥ 70%. The cytotoxic compounds were further filtered out if cell count ≤ 50% of neg ctrl from the imaging data. The final hit list included 109 compounds representing 2.2% hit rate. (E) Overview of hit numbers from each assay/analysis: Cell Painting linear predictive model, Cell Painting supervised PCA, and CXCL10 reduction assay.

In addition to CXCL10 protein as a readout for efficacy we also used the Cell Painting assay to serve as a morphological readout of cellular fibrosis. Morphologies of CCD-18co cells treated with different profibrotic stimuli were visually distinct (S. Figure 4A) and this translated to cellular features that yielded equally distinct principal component analysis (PCA) plots (S. Figure 4B). Interestingly, the Cell Painting PCA plot strongly resembled the transcriptomic PCA plot (S. Figure 4C), suggesting CCD-18co cellular morphology might be tightly correlated to gene expression and subsequent biological activities.

**Figure 4.**
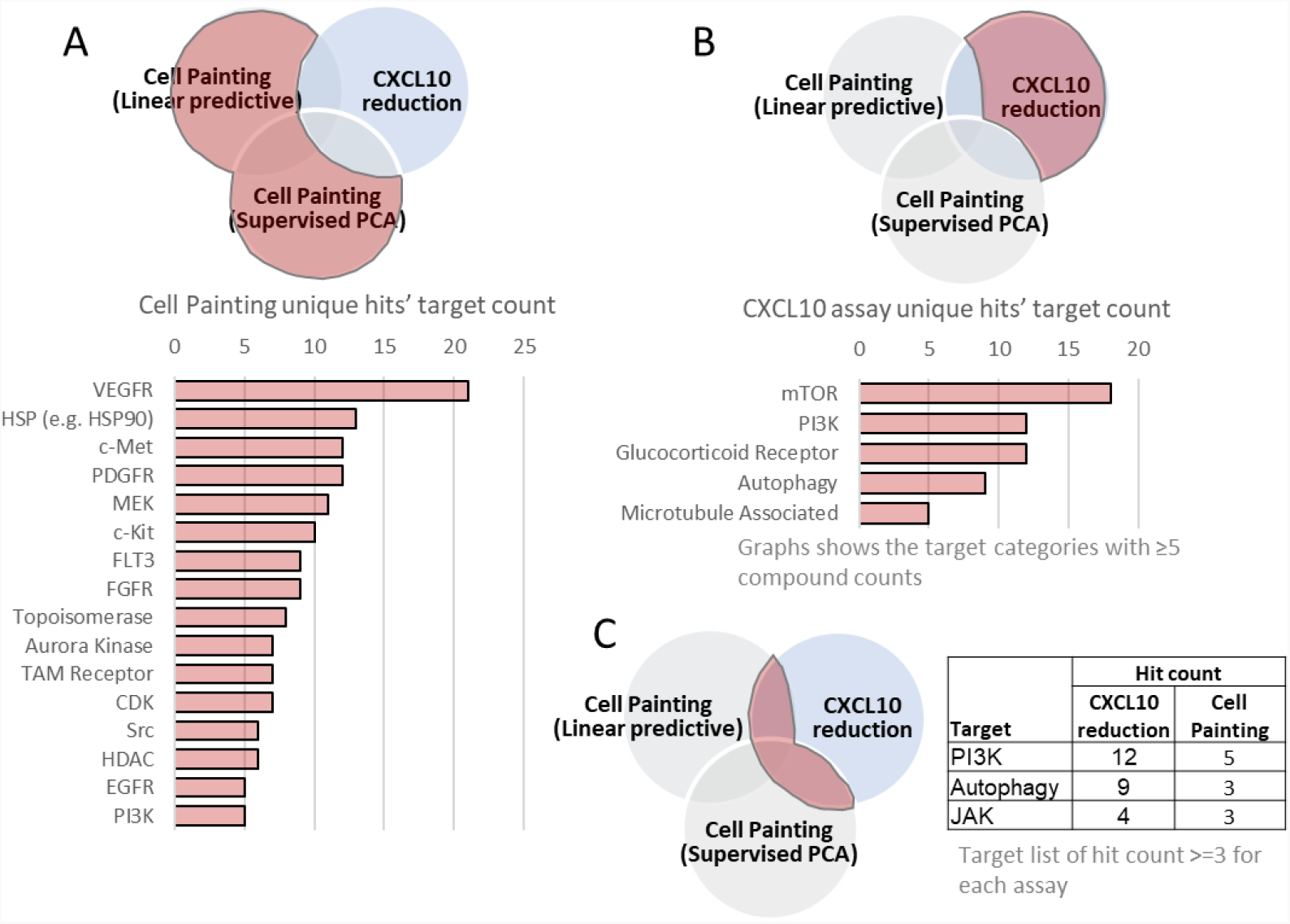
Hit category analysis of Cell Painting and CXCL10 reduction assays. (A) Hit number of target categories for linear predictive model and supervised PCA analysis of Cell Painting. Bar chart shows the target categories with ≥5 compounds in each. (B) Hit number of target categories for CXCL10 reduction assay. Bar chart shows the target categories with ≥5 compounds in each. (C) Hit number of target categories for the overlapping hits between CXCL10 reduction assay and Cell Painting assays. Table shows target categories with ≥3 compounds in each assay.

### Automated high throughput chemogenomic library screen to identify targeted perturbagens of intestinal fibrosis

To comprehensively profile diverse biological and functional space (Figure 2A), we sourced two small molecule libraries totaling 4871 compounds annotated with either their reported targets and/or mechanisms of action and have been either tested in clinical trials or approved by the FDA (Selleckchem; S. Figure 5A). The molecular weight and ALogP of these compounds were within the standard range for “drug-like” molecules (S. Figure 5B)^*17*^.

**Figure 5.**
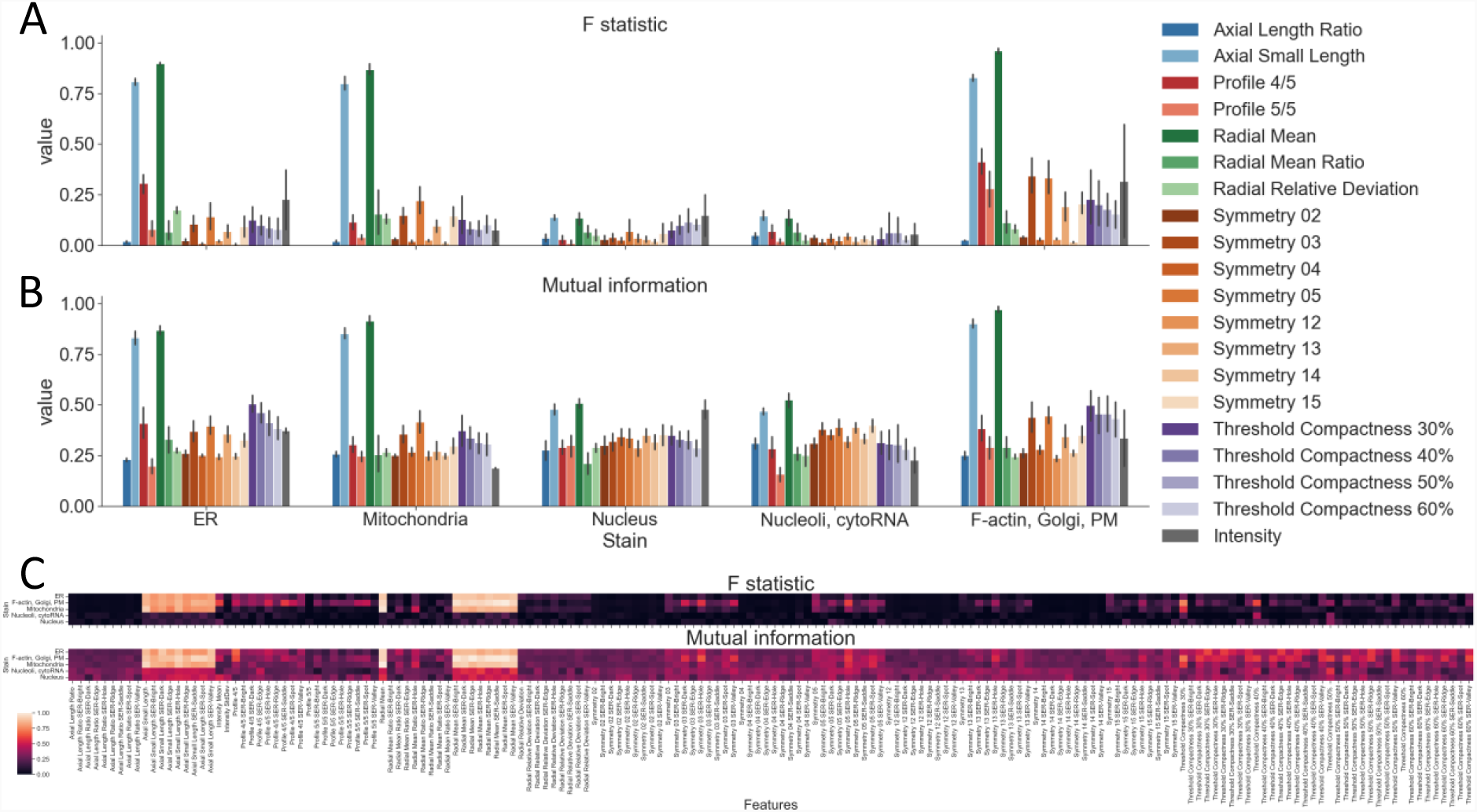
Correlation analysis of morphological features with CXCL10 level. (A and B) F statistic (which shows the linear dependency), and mutual information (which shows any type of dependency, including linear dependency), between cellular feature subcategories and the CXCL10 level. (C) Heat map of top highly correlated features of each subcategory with CXCL10.

For the primary high throughput screening assay, 1,200 CCD-18co cells/well were plated on the first day, followed by compounds and controls after 24 hours (S. Figure 5C). Each compound was tested at 3 μM in biological triplicate. 1 ng/mL anti-TNFα antibody Adalimumab was used as the positive control, because Adalimumab was able to effectively suppress TNFα signaling in the CXCL10 assay (as well as in the Cell Painting assay, as discussed later, S. Figure 6). Cells were then treated with 10 ng/mL TNFα on the third day for 48 hours, after which time the cell culture supernatant were collected for CXCL10 protein quantitation using a Homogeneous Time Resolved Fluorescence (HTRF) assay.

**Figure 6.**
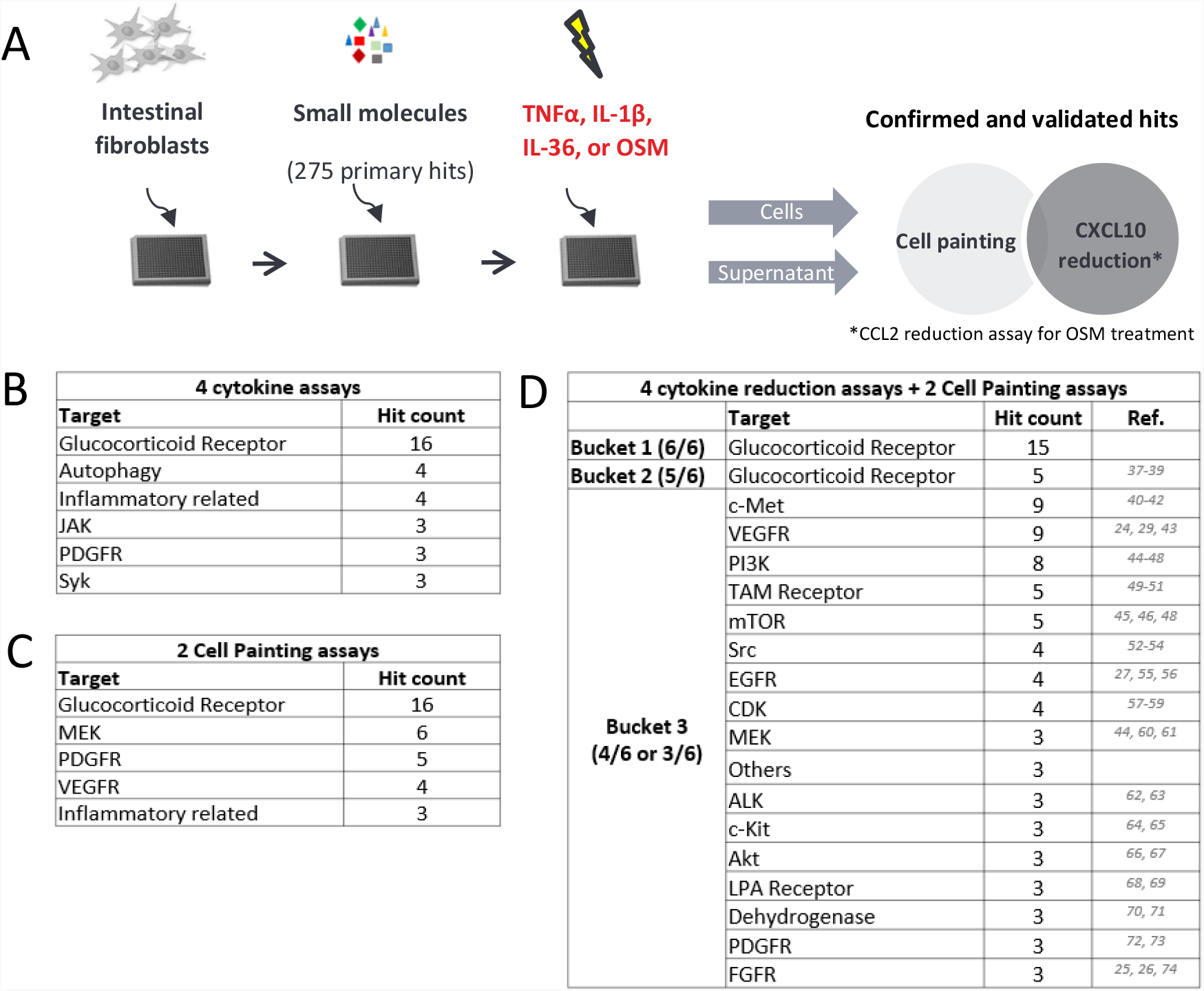
Hit confirmation and validation assay workflow and hit categories. (A) Hit confirmation and validation experimental workflow. Intestinal myofibroblasts CCD-18co cells were seeded in 384 well plates. After overnight incubation, 275 small molecule hits from the primary screen, positive, or negative controls were dispensed into the wells. After 24-hour incubation, cells were treated with TNFα, IL-1β, IL-36 or OSM. After 48-hour stimulation, the cell culture supernatants were collected to assess CXCL10 levels for TNFα, IL-1β, IL-36 treated samples and CCL2 levels for OSM treated samples. The cells were stained with Cell Painting dyes for morphological profiling. There was an insufficient assay window between positive and negative controls for Cell Painting results from IL-36 and OSM treated samples, so Cell Painting results were not produced for these stimuli. (B) The top target categories across the four cytokine reduction assays. Table shows target categories with ≥3 compounds for each assay. (C) The top target categories for the Cell Painting results of TNFα and IL-1β stimulation. (D) The top target categories for all six assay results. The results were further bucketed into three categories. Bucket 1 includes compounds that had effect in all 6 assays. Bucket 2 includes compounds that had effect in 5 out of 6 assays. Bucket 3 includes compounds that had effect in 3 or 4 out of 6 assays. Table only shows target categories with ≥3 compounds in each.

For the Cell Painting assay, the cells from the exact same samples were stained with Cell Painting dyes followed by high content image acquisition and analysis. The assay includes six fluorescent dyes to highlight different organelles of CCD-18co cells, including MitoTracker™ Deep Red FM for mitochondria, Concanavalin A-Alexa 488 for endoplasmic reticulum, SYTO™ 14 for nucleoli and cytoplasmic RNA, WGA-Alexa 555 and phalloidin-Alexa 568 for F-actin cytoskeleton, Golgi and plasma membrane, Hoechst 33342 for nucleus^*18*^ (Figure 2B). High content images were captured and cellular morphological features were extracted, then analyzed using a dimensionality reduction method. Compounds that clustered around the positive controls were categorized as Cell Painting hits (Figure 2C). For dimensionality reduction, we used either a supervised PCA or a linear predictive model. For both methods, the medians of positive controls and negative controls were normalized to 0 and 1, respectively. Compounds were then binned into positive or negative bins depending on the projection scores (Materials and methods, Figure 3A, left). Compounds positive for >2 out of 3 replicates in the positive bins and projection scores within the range of Average(pos ctrl) ± 3 x S.D. (standard deviations) were picked as preliminary hits. Compounds exhibiting cytotoxic profiles were then further filtered based on cell count.

In total, 160 and 152 compounds were picked as hits from supervised PCA and linear predictive models of the Cell Painting data, respectively (Figure 3A, right). There were 100 hits that overlapped between both models for Cell Painting analysis (Figure 3A, right), suggesting the two analytical methods yielded mainly convergent results. In addition, we assessed three other metrics for picking Cell Painting hits; namely using the top 50 features, top 5 features, or top 3 features per channel that separate positive and negative controls, though the hits and targets that were identified were mostly similar (S. Figure 7). To determine whether these cellular features correlate with their biological functions, we projected the cellular features of the targets that were most distinct from the negative controls onto a two-dimensional t-SNE map (Figure 3B). This map showed that some co-annotated compounds form coherent clusters (e.g. MEK and HSP) in phenotypic space whereas other do not (e.g. Bcl-2, FAK, CRM1, DNA-PK).

**Figure 7.**
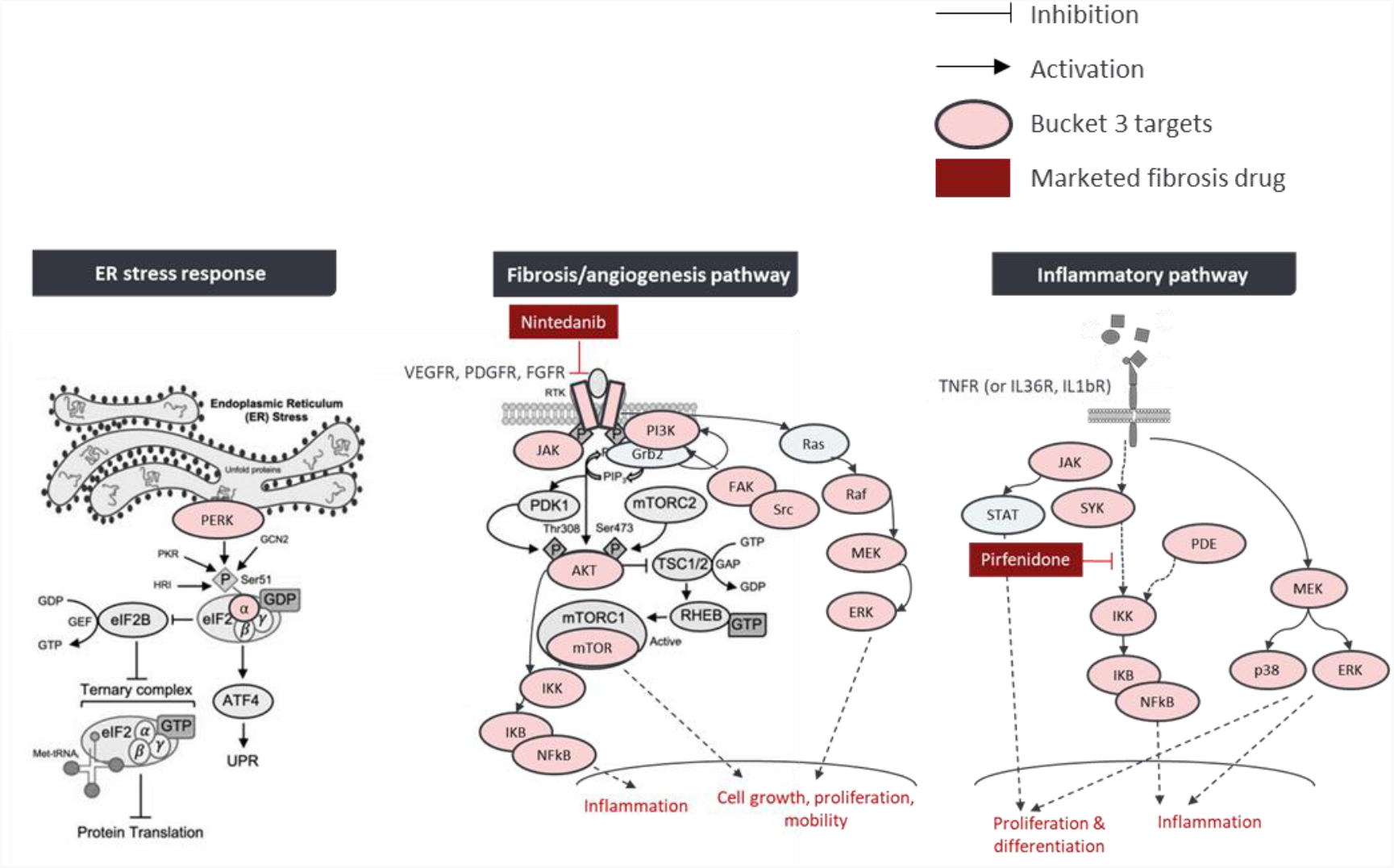
Major pathways of the bucket 3 compound targets. Three major pathways, including ER stress response, fibrosis/angiogenesis pathway, and inflammatory pathway were identified by analyzing the targets of bucket 3 compounds. Pink bubbles show the targets that were identified in the bucket 3 compounds. Gray bubbles show other intermediate targets in the pathway. Nintedanib, a marketed drug for idiopathic pulmonary fibrosis, was identified as a hit in the screen. The screen also identified inflammatory pathway targets through which pirfenidone, another marketed drug for idiopathic pulmonary fibrosis, exerts its effect.

For the CXCL10 assay, luminescence intensities of positive and negative controls of each plate were fit on a 0 to 1 scale, and were then normalized for their percent inhibition, with the mean of positive control being 100% and the mean of negative control being 0% (Figure 3C). The strictly standardized mean difference (SSMD) was used to measure the effect size and gauge the assay quality^*19*^. Plates with SSMD>1.28 (the SSMD quality cutoff) then proceeded to hit selection. Compounds positive for >2 out of 3 replicates with CXCL10 inhibition > 70% were identified as preliminary hits, then filtered by eliminating cytotoxic compounds (dependent on cell count). After applying this gating strategy, 109 compounds were identified, resulting in a 2.2% hit rate for the CXCL10 screen (Figure 3D).

Surprisingly, there were only 8 hits that overlapped between the Cell Painting linear predictive model and the CXCL10 assay, and only 9 hits that overlapped between the Cell Painting supervised PCA model and the CXCL10 assay. Only 5 hits overlapped among all three methods. In the end, after removing duplicate compounds, 275 unique hits from either Cell Painting or CXCL10 assay were advanced for further confirmation and validation (Figure 3E).

### Target discovery through the integration of cytokine biomarker and morphological profiles

It was intriguing that the CXCL10 assay and the Cell Painting assay identified vastly different pools of hit compounds. For hit compounds that were unique to Cell Painting, the top targets included VEGFR, HSP, c-Met and PDGFR, MEK, c-Kit, FLT3 and FGFR (Figure 4A); while hits that were unique to the CXCL10 assay included the targets mTOR, PI3K, glucocorticoid receptor and several components of the autophagy and microtubule pathways (Figure 4B). For the hits that were shared between the two assays, the top targets included PI3K, autophagy and JAK (Figure 4C).

In several contexts, image-based profiles have proven to have predictive ability for other assays^*20*^. We wondered whether any particular cellular morphology features from the Cell Painting assay can be used to predict CCD-18co cells’ response to TNFα, in terms of secreting CXCL10. We studied the statistical dependence between CXCL10 levels and each of the 860 individual cellular features. We found that a few subcategories of cellular features including Axial Small Length (the length of the cell’s shorter axis in pixel units) and Radial Mean (the mean object radius based on the intensity values weighted by the distance from the mass center) from the ER, mitochondria and F-actin, Golgi and PM channels had strong relationship with CXCL10, as F statistic values which capture the linear dependency between features and the CXCL10 outcome were 0.73 on average (Figure 5A), and the mutual information (MI) values which capture any type of dependency were 0.76 on average (Figure 5B). To zoom in the subcategories and examine which particular features had the strongest statistical dependency with CXCL10 level, we identified a few Radial Mean features, including Edge, Ridge, and Spot of the SER texture analysis in the F-actin, Golgi, plasma membrane channel have nearly perfect statistical dependency with CXCL10 level (e.g., Radial Mean SER-Spot has F-statistic of 1.0 and Radial Mean SER-Edge has MI of 1.0) (Figure 5C), indicating that these features have strong dependency with CXCL10 and can be used to predict CXCL10 level.

### Target validation using pro-fibrotic stimuli-treated cell models

To further characterize hit compounds according to their ability to ameliorate fibrosis from pro-fibrotic stimuli other than TNFα, we profiled the 275 unique hit compounds at three doses (3 μM, 0.6 μM and 0.125 μM) in assays with different stimuli (IL-1β, IL-36 or OSM) in addition to TNFα (Figure 6A). The CXCL10 assay was used for TNFα-, IL-1β-and IL-36-treated cells, while a CCL2 assay conducted 2 hours post OSM treatment was used for OSM-treated cells, because CCL2 (S. Figure 8) but not CXCL10 (S. Figure 2) is a functional biomarker for OSM stimulation. The Cell Painting assay was only used for TNFα and IL-1β stimulation, as there were not viable assay windows for cells treated with either IL-36 or OSM (S. Figure 4B), leaving four cytokine assay and two Cell Painting assay results available for analysis.

The TNFα-stimulated reconfirmation screen of 275 unique hit compounds yielded a 51% reconfirmation rate for reducing CXCL10 expression/secretion and a 47% reconfirmation rate for Cell Painting, suggesting robustness of the primary screening assays. Using a combinatory approach to examine the target categories, we pooled the four cytokine stimulation results and identified glucocorticoid receptor as the top target with 16 hits. This was followed by autophagy, inflammatory-related mechanisms, JAK, PDGFR and SYK (Figure 6B). The two Cell Painting reconfirmation assays (TNFα and IL-1β) similarly showed glucocorticoid receptor to be the top target, followed by MEK, PDGFR, VEGFR and inflammatory-related mechanisms (Figure 6C).

When considering all six compound lists, the hits were binned into three buckets depending on the number of assays they were identified as hits. Bucket one included compounds that were picked as hits in six out of six assays. All hits in this bucket were glucocorticoid receptor modulators (steroids). Bucket two included compounds that were picked as hits in five out of six assays and similarly, all hits in bucket two were mainly glucocorticoid receptor modulators. Bucket three included compounds that were identified as hits in three or four out of six assays and this bucket represented the largest variety of biological functions with different mechanisms of action (Figure 6D).

To understand these targets in the context of signaling pathways, we mined the literature and identified any associations between targets in bucket three and intestinal fibrosis. Overall, three main pathways were identified: ER stress response, fibrosis/angiogenesis and inflammation (Figure 7). All three pathways were shown to play a role in tissue fibrosis^*14, 21-28*^. Interestingly, we identified and confirmed both Nintedanib (targets PDGFR, VEGFR and FGFR) and Pirfenidone (targets NF-kB), which were approved drugs for treating idiopathic pulmonary fibrosis (IPF)^*28, 29*^, as potent antagonists of myofibroblast activation^*30*^ (Figure 7). These data suggest that the small molecules, targets and signaling pathways identified through our multi-parametric biomarker and cellular feature profiling approach were physiologically-and clinically-relevant. Further, this screening platform was able to identify molecules from a wide spectrum of mechanisms of action.

## Discussion

IBD-associated intestinal fibrosis represents a highly invasive and deleterious disease that currently has no approved pharmacological intervention. In order to address this as well as the gap in clinically reliable rodent preclinical models (ref for this statement) we developed a clinically relevant humanized intestinal fibrosis model composed of TNFα-activated colon fibroblasts. In order to leverage large collections of small molecules for therapeutic profiling efforts, we miniaturized the human IBD fibrosis model to accommodate a scalable phenotypic screening platform for fully-automated drug discovery. Employing transcriptomics as a surrogate characteristic for comparing our CCD-18co *in vitro* model to IBD patient biopsies, we identified several distinct transcriptional clusters corresponding to different pro-inflammatory cytokine stimuli. Although TGFβ treatment of CCD-18co cells produced a canonical gene expression profile that overlapped with myofibroblast components of patient biopsies, TNFα stimulation yielded a fibrotic phenotype most consistent with the IAF population of IBD patient samples.

As intestinal fibrosis is a result of a complex interplay of immune-mediated inflammatory processes as well as modulation of pro-inflammatory cytokine-mediated signaling pathways, our screening platform required a sophisticated series of assay readouts to account for these polyetiological causes. We first chose to use CXCL10 (IP10) as a robust biomarker for fibrosis due to its well-characterized association with intestinal fibrotic pathology and because compared to other biomarkers both its mRNA and protein levels were significantly promoted by multiple pro-fibrotic stimuli (S. Figure 2). However, to fully assess changes in the fibrotic morphological phenotype, we developed an unbiased image-based profiling technique called Cell Painting. Although Cell Painting has not been widely adapted in the drug discovery industry as a phenotypic readout for efficacy, its scalable ease of use as well as its ability to quantitate changes in thousands of cellular features make it an ideal method for studying complex biology such as intestinal fibrosis. Cell Painting produces vast morphological information as a collection of extracted cellular features, but by integrating artificial intelligence analytical methods, such as machine learning, we can mine these data to reveal important biological activities of potentially therapeutic small molecules^*12*^. For example, we found that the relative positions of pro-fibrotic stimuli-treated clusters to vehicle controls in Cell Painting PCA plots were similar to those from RNA-seq PCA plots, suggesting transcriptome profiles and related biological activities strongly correlate with cellular morphological profiles. We also examined whether any specific cellular features were highly correlated with CXCL10 level, because these features may potentially be used as sentinel readouts for CXCL10 in future studies. We identified several subcategories of features, such as Axial Small Length and Radial Mean in ER, mitochondria and F-actin, Golgi and plasma membrane channels that had high correlations with CXCL10 level (Figure 5).

Surprisingly, we observed divergent hit distribution profiles between the CXCL10 and Cell Painting assay readouts. While the CXCL10 readout identified well-characterized regulators of fibrosis such as mTOR and glucocorticoid receptor, the targets identified through the Cell Painting readout were mechanistically more diverse (e.g. VEGFR, PDGFR, FGFR, c-Met, c-Kit and MEK) and included such cellular processes as fibrosis, tissue plasticity and remodeling, and angiogenesis. In short, the CXCL10 assay conferred a confidence metric to the biological relevance of our assay platform by identifying several steroid molecules as alleviators of the fibrotic phenotype but the Cell Painting assay was able to reveal a wider range of potentially-actionable (and novel) mediators of intestinal fibrosis pathology.

As our collective understanding of the causes and mediators of disease biology increase, so must our ability to interrogate those causes to discover the next generation of small molecule therapeutics. A complex image-based profiling technique like Cell Painting integrated with state-of-the-art machine learning algorithms to translate thousands of cellular features into disease-relevant targets and pathways may represent a giant leap forward in industrialized drug discovery. Although it may be unlikely that image-based profiling will completely replace conventional biochemical, transcriptional or proteomic profiling methods, when incorporated into exploratory phases of the drug discovery pipeline, Cell Painting may accelerate identification of novel therapeutics and expand the targeting space of polyetiological and poorly-understood diseases like intestinal fibrosis.

## Materials and methods

### Single Cell RNA-seq

CCD-18co cells were cultured as previously described, and were treated with vehicle, 10 ng/mL OSM, 10 ng/mL TNFα, 10 ng/mL IL-1β, 100 ng/mL TL1A, 200 ng/mL IL-36 or 10 ng/mL TGFβ for 16 hours. Cells were detached from plastic culture media using Trypsin EDTA and manually counted. 10,000 cells of each sample were loaded in individual lanes of a 10X Genomics Next-GEM chip (10X Genomics, cat #: 1000127). Libraries were prepared using the 10X Genomics Next-GEM 3’ GEM kit (10X Genomics, cat #: 1000121). Library quality control was performed using the Agilent Tapestation D1000 and D5000 tapes (Agilent, part #: 5592 and 5584). Sequencing was performed externally at a vendor (Novogene Inc.) at a depth of 50,000 reads per cell.

### Analysis of CCD18Co scRNA-Seq data

Cell Ranger mkfastq was used to demultiplex raw sequencing reads, and Cell Ranger count was used to align reads to Human GRCh38 transcriptome, and generate gene-cell expression count matrices. Expression matrices were filtered to remove low quality cells with less than 200 genes detected or more than 0.25% mitochondrial mRNAs. Cell filtering resulted on an average of ∼2,400 cells per condition and a total number of 16,750 cells in the integrated dataset. Seurat v3[Butler et al., 2018] workflow was employed to perform normalization, detection of variable genes, dimensionality reduction, and graph-based clustering with Louvain algorithm. Upon log-normalization and scaling of gene expression, variable genes were identified using the vst method and then subjected to principal component analysis (PCA). The number of principal components (PCs) used for nonlinear dimensional reduction analysis (UMAP) was chosen according to the PCElbowPlot function and JackStrawPlot function. For cell clustering, FindClusters method was parameterized with different resolutions to optimize cluster granularity. Sample integration was performed with the IntegrateData function using anchors set by FindIntegrationAnchors. Cells with high mitochondrial RNA expression (greater than 5% of total cell reads) were excluded from downstream analysis. FindAllMarkers function in Seurat was utilized to detect top gene markers per cluster with the default Wilcoxon rank-sum test setting. The results of FindAllMarkers were subjected to functional pathway analysis. Upregulated gene markers per cluster with logFC>0.4 and adjusted p-value <0.05 were subjected to functional enrichment with Ingenuity Pathway Analysis (IPA) (Qiagen Inc.). IPA canonical pathways and upstream transcriptional regulators were predicted per cluster and then integrated via the Comparison Analysis function.

### CCD-18co cell culture and screening assay

A large batch of CCD-18co cells were purchased from ATCC (Cat #: CRL-1459), and were cultured with ATCC-formulated Eagle’s Minimum Essential Medium (Cat #: 30-2003) with 10% fetal bovine serum (FBS) as recommended by ATCC. The cells earlier than passage-6 were used in experiments. The chemogenomic screen library was assembled with two screening libraries from Selleckchem (cat #: L1100 and L3800). The high throughput screen was conducted with an automation platform consisting of multiple instruments (S. table 1), which were operated by Green Button Go software (Biosero). For pro-fibrotic stimuli treatments, if not specified otherwise, the concentrations of stimuli used to treat CCD-18co cells were as follow: 10 ng/mL OSM, 10 ng/mL TNFα, 10 ng/mL IL-1β, 100 ng/mL TL1A, 200 ng/mL IL-36 and 10 ng/mL TGFβ.

For the primary screen, 1,200 cells/16 μL/well of CCD-18co cells were seeded in CellCarrier-384 Ultra microplates (Perkin Elmer, Cat #: 6057302) with an EL406 combination microplate washer dispenser (EL406, Biotek). After overnight incubation at 37 °C, 37.5 nL/well (for 2 mM stock) or 7.5 nL/well (for 10 mM stock) small molecules were dispensed by ATS GEN5 (EDC Biosystems). The wells were then backfilled with 10 μL/well cell culture medium with an EL406 (except for column 12). Column 12 was filled with 10 μL/well adalimumab (final concentration 1 nM) for positive control wells or 10 μL/well medium for negative controls. After overnight incubation, each well was then filled with 25 μL/well recombinant human TNFα (R&D systems, cat #: 210-TA-100) diluted in cell culture medium (final concentration 10 ng/mL) with an EL406. After 48-hour stimulation, 4 μL/well of cell culture supernatant was taken out and dispensed into OptiPlate-384 microplates (PerkinElmer, cat #: 6007290) that was pre-filled by 12 μL/well PBS by an EL406. CXCL10 HTRF assay antibodies were prepared according to the manufacturer’s recommendation (Cisbio, cat #: 62HCX10PEH). 4 μL of the mixture of equal part of diluted CXCL10 Eu Cryptate antibody and CXCL10 d2 antibody was added to PBS diluted supernatant in OptiPlate-384 microplates by a Bravo liquid handler. The OptiPlate-384 microplate was then sealed by a PlateLoc thermal microplate sealer (Agilent) and shaken at 1000 rpm for 30 seconds on a BioShake 5000 elm shaker (QInstrument) followed by centrifugation at 500xg for 30 seconds on a Microplate centrifuge (Agilent). The OptiPlate-384 plate was incubated at room temperature in a Biosolutions AmbiStore™ D (HighRes) for 2 hours before the HTRF signal was measured on an EnVision® HTS plate reader (PerkinElmer). For OSM-induced CCL2 assay in hit validation step, the cell supernatant was collected after 2-hour treatment with OSM. The experimental procedure was similar to CXCL10 assay, except the signal was detected using a CCL2 kit (Cisbio, Cat #: 62HCCL2PEG). The OSM treated cells were then returned back to the incubator, and the Cell Painting assay was carried out after 48 hours post OSM treatment. After supernatant being transferred to the OptiPlate-384 microplates, the cells in CellCarrier-384 Ultra microplates were treated with 25 μ L of 0.71 μM MitoTracker™ Deep Red FM (ThermoFisher, cat #: M22426) that was prepared in cell culture medium from 1 mM DMSO stock (final concentration of MitoTracker™ Deep Red FM is 0.25 μM). MitoTracker™ Deep Red FM was dispensed by an EL406. After incubation with MitoTracker™ Deep Red FM for 30 minutes at 37 °C, the cells were then fixed with 4% paraformaldehyde for 10 minutes. Paraformaldehyde was dispensed by an EL406. The CellCarrier-384 Ultra microplates were then washed with PBS twice on a Blue®Washer, and further stained with a cocktail of 15 μL/well including 2 μg/mL Hoechst 33342 (Invitrogen, cat #: H3570), 100 μg/mL Concanavalin A-Alexa 488 (Invitrogen, cat #: C11252), 3 μM SYTO™ 14 (Invitrogen, cat #: S7576), 5 μg/mL WGA-Alexa 555 (Invitrogen, cat #: W32464), 2.5 u/mL phalloidin-Alexa 568 (Invitrogen, cat #: A12380) for 30 minutes at room temperature. The dye cocktail was dispensed by a Blue®Washer. After incubation, the plates were washed with 1X PBS three times. The cells were then imaged with an Operetta CLS High Content Analysis System (PerkinElmer).

MitoTracker™ Deep Red FM was reconstituted with DMSO and was always prepared fresh from lyophilized stock. Concanavalin A-Alexa 488 was diluted with 0.1 M sodium bicarbonate and WGA-Alexa 555 was diluted with ddH2O, and unused stock of both could be frozen in -20 °C up to 1 month. Phalloidin-Alexa 568 was reconstituted with DMSO. Both phalloidin-Alexa 568 and SYTO™ 14 could be frozen in -20 °C up to 1 year. Hoechst 33342 was stored at 4 °C.

### Cell Painting high content imaging, feature extraction and high dimensionality data analysis

Cells stained with Cell Painting dyes on CellCarrier-384 Ultra microplates were imaged with an Operetta CLS High Content Analysis System. The details of the Operetta CLS channels and stains imaged in the Cell Painting assay are shown below.

| Dye | Filter (Excitation) | Filter (Emission) | Organelle or cellular component |
| --- | --- | --- | --- |
| Hoechst 33342 | 355-385 nm | 430-500 nm | Nucleus |
| Concanavalin A-Alexa 488 | 460-490 nm | 500-550 nm | Endoplasmic reticulum |
| SYTO™ 14 | 490-515 nm | 525-580 nm | Nucleoli, cytoplasmic RNA |
| WGA-Alexa 555 and phalloidin-Alexa 568 | 530-560 nm | 600-640 nm | F-actin cytoskeleton, Golgi, plasma membrane |
| MitoTracker™ Deep Red FM | 615-645 nm | 685-760 nm | Mitochondria |

20x water objective with non-confocal mode was used. Binning value of 2 was chosen to boost camera frame rate and dynamic range. The middle 9 fields of view were imaged for each well. The exposure time and power were adjusted to ensure the max of signal level under auto contrast for each channel was between the range of 8,000 – 15,000.

For Cell Painting feature extraction, Harmony® PhenoLOGIC™ software was used. The basic flatfield correction and brightfield correction were applied to the images. Hoechst 33342 channel was used to identify nuclei, followed by using concanavalin A-Alexa 488 channel to find cytoplasm. Border objects of each image were removed to ensure the cells with complete image were used for feature extraction. Properties of ell features were then quantified, including fluorescnece intensities within different cell regions, basic morphological features (cell area, roundness, length, width and width/length ratio), advanced morphological features (<u>S</u>ymmetry, <u>T</u>hreshold compactness, <u>A</u>xial, <u>R</u>adial and profile, or STAR properties), cell texture features (<u>S</u>pot, Hole, <u>E</u>dge, <u>R</u>idge, Valley, Saddle, Bright and Dark, or SER properties)^*31*^. In total, there were 860 cellular features being extracted.

The data including all extracted cellular features was imported to TIBCO® SpotFire® Signals VitroVivo (PerkinElmer) for analysis. Within the software, Editable Data Grid application was used to associate cellular feature data file with the compound transfer log file to annotate compounds. Grid Plate Editor application was used to designate the plate layout. High Content Profiler was then used to perform advanced feature normalization, selection, classification, profiling and hits selection. The compounds with score in the range of Average(neg ctrl) ± 3 S.D. were selected. If they appear 2 times among the tested triplicates, they were picked as hits.

In parallel, the Cell Painting data was analyzed with a linear predictive model. Briefly, the positive and negative controls were used as a training set to train a predictive model. Top features were ranked using area under the ROC curve. Multiple testing adjusted p-values were also calculated for each feature using a random permutation method. The top features that were highly predictive and could separate positive and negative controls were used to rank compounds. The compounds with scores in the range of Average(neg ctrl) ± 1 S.D. and appeared at least 2 times were picked as hits.

### Cell Painting data projection score calculation

Projection score is a mathematical formalization of a heuristic, which predict sample’s positivity depending on whether the feature values are closer to the positive controls and the negative controls. Projection score is calculated as

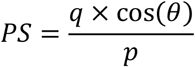

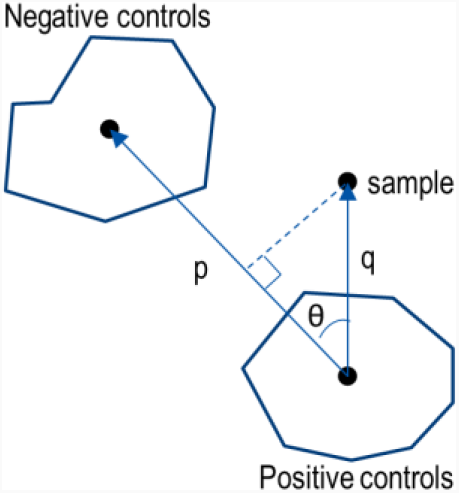

Where ***p*** is the distance between the center of positive and negative controls, ***q*** is the distance between sample and the center of negative controls, and *θ*is the angel between side ***q*** and side ***p***.

### CCD-18co cell hit confirmation and validation hit picking strategy

For cytokine (CXCL10 and CCL2) assay hit picking, the assay plates first passed SSMD>1.28 QC criteria. Compounds’ cytotoxicity effects were assessed by cell count from the imaging assay. If three or more cell count data points out of the six data points (three doses in replicates) of a compound show < 50% cell viability compared to the negative control, this compound was then considered toxic and filtered out. The area under the curve (AUC) of six normalized CXCL10 percent inhibition data points of each compound was then calculated for each stimulus. The compounds were ranked according to the values of AUC. For Cell Painting assay hit picking, after elimination of the cytotoxic compounds, the projection scores of Average(pos ctrl) ± 3 x S.D. were selected. If a compound had three or more data points out of the six data points that met this criterion, it was confirmed as a hit.

### Visualization of primary screen Cell Painting dataset with t-SNE

To generate t-SNE projection of Cell Painting morphological feature profiles, compounds that have target annotations were selected and per-compound feature profiles were aggregated by calculating the mean of each feature across compound replicates. Per-target feature profiles were then aggregated by calculating the mean of each feature across all compounds annotated with this target. Per-target feature profiles were then rank-ordered by decreasing Euclidean distance from the mean feature profile of negative controls. Finally, the top-20 most distinct targets were selected and the targets that were associated with 3 or less compounds were filtered out, obtaining the final set of 13 targets.

The two-dimensional t-SNE embedding was generated using the scikit-learn Python library with random initialization^*32*^, perplexity of 20, early exaggeration of 30, automatic learning rate and squared cosine distance as a metric. Targets that formed coherent clusters were highlighted using kernel density estimation with empirically chosen thresholds. It should be noted that cluster sizes and inter-cluster distances should be interpreted with care when using t-SNE^*33*^. The t-SNE visualization was built using matplotlib^*34*^ and seaborn^*35*^ Python libraries.

### CXCL10 and cellular feature statistical dependence analysis

To assess the correlation between CXCL10 level and cellular features extracted from Cell Painting imaging data, we first filtered out those compounds that demonstrated cytotoxic effects. We then selected those compounds that were present in both CXCL10 and imaging assays and aggregated per-compound imaging feature profiles by calculating the mean of each feature across compound replicates. Then, for each cellular morphological feature, we computed two metrics of relatedness to CXCL10 level: F-test statistic and Mutual information (MI). The method based on the F-test statistic estimates the degree of linear dependency between the feature and the outcome, while the mutual information method can capture any kind of statistical dependency. We plotted obtained results either as individual data points (Figure 5B) or as a bar plot that aggregates features by their type (Figure 5A), with error bars showing a bootstrap-estimated 95% confidence interval.

Computations of the F-test statistic and mutual information were performed using the scikit-learn^*32*^ and pandas^*36*^ Python libraries. Visualizations were built using matplotlib^*34*^ and seaborn^*35*^ Python libraries.

### CRISPR/Cas9, mismatch detection and T7E1 assays

Lentiviral particles of inducible Cas9 nuclease with hEF1a promoter (Horizon discovery, cat #: VCAS11227) were transduced into CCD-18co cells with MOI=0.5 according to manufacturer’s instruction. The Cas9 expressing cells were selected under 2 μg/mL blastcidin. The cells were added 10 ng/mL doxycycline to induce Cas9 expression. The Cas9 expression could be readily detected by Western blot after 24 hours. CRISPR gene knockout kits or individual gRNA for targeting *TNFRSF1A* or *TNFRSF1B* were ordered from Synthego. For gRNA transfection, 0.05 μL Lipofectamine™ RNAiMAX (ThermoFisher, cat#: 13778075) mixed with 5 μL opti-MEM was dispensed into each well of CellCarrier-384 Ultra microplates, followed by dispensing 2 pmol gRNA diluted in 5 μL opti-MEM. The RNAiMax and gRNA were then mixed by shaking the plate on a microplate shaker for 1 minute at 800 rpm. The plate was then centrifuged 1 minute at 500xg and incubated at room temperature for 20 minutes. After incubation, 1,200 cells/well with 10 ng/mL doxycycline were seeded in CellCarrier-384 Ultra microplates with RNAiMax and gRNA mixture. After 72 hours, the cells were collected for T7E1 assay.

For mismatch detection and T7E1 assay, the cells were lysed in 20 μL of 1x Phusion high-fidelity buffer (Thermofisher, cat #: F-518L) with 1 mg/mL proteinase K (Thermofisher, cat #: EO0492) and 0.5 mg/mL RNase A (cat #: EN0531). The plate was then incubated for 15 – 30 minutes at 56 °C, followed by deactivation for 5 minutes at 96 °C. Briefly centrifuge plate to collect liquid at bottom of wells. A 50 μL PCR reaction with the following condition was set up: 1X Phusion High-Fidelity buffer (cat #: F-549S), 500 nM forward primer, 500 nM reward primer, 200 μM each dNTPs, 0.04 U/μL Phusion hot start II highfidelity DNA polymerase (cat #: F-549S), and 5 μL cell lysate. The thermal cycling condition is as shown below:

| Cycle step | Temperature (°C) | Time | Cycle(s) |
| --- | --- | --- | --- |
| Initial denaturation | 98 | 3 min | 1 |
| Denature | 98 | 10 s | 10 |
| Touchdown annealing | 72 – 1/cycle | 15 s |  |
| Extension | 72 | 30 s |  |
| Denature | 98 | 10 s | 25 |
| Annealing | 62 | 15 s |  |
| Extension | 72 | 30 s |  |
| Final extension | 72 | 10 min | 1 |

PCR samples were heated to 95 °C for 10 minutes and then slowly (>15 minutes) cool to room temperature. 15 µL mismatch detection assay using T7E1 was then set up. The reaction includes PCR from gDNA (300 – 500 ng/reaction), 1X NEBuffer 2 (NEB, cat #: B7002S), and 0.33 U/µL T7 endonuclease I (NEB, Cat#: M0302S). The mixture was then incubated for 25 minutes at 37 °C. The entire reaction was then loaded on 2% agarose gel to determine gRNA cleavage.

### Olink Target 96 Inflammation assay

CCD-18co cells were treated with vehicle, 10 ng/mL OSM, 10 ng/mL TNFα, 10 ng/mL IL-1β, 100 ng/mL TL1A, 200 ng/mL IL-36 or 10 ng/mL TGFβ for 16 or 48 hours. The cell culture supernatant samples (n=5 for each treatment) were collected and sent to Olink for the Olink Target 96 Inflammation assay. The samples were tested in neat or in 1:10 dilution. In total, 92 immune-related secretory biomarker protein levels were assessed. The results were presented in Normalized Protein Expression (NPX) value, which is an arbitrary unit on a Log2 scale. The linear portion of the NPX vs. concentration plot was used for calculation. The NPX values were further normalized to the vehicle treated control group. The heat map shows the average of the protein concentration of n=5. In parallel, the mRNA samples of the cells that were treated with various stimuli for 16 hours were sequenced. The matching gene expression levels of the 92 proteins that were assessed in the Olink assay were analyzed. The FPKM levels for each treatment was normalized to the vehicle control group. The heat map shows the average of the gene expression level of n=3.

### RNA-seq

CCD-18co cells were treated with various stimuli for 16 hours in the same way as described above in *Olink Target 96 Inflammation assay* section. For mRNA-seq, the raw reads were aligned to the transcriptome using STAR (version 2.6.0)/RSEM (version 1.2.25) with default parameters, with a custom human GRCh38 transcriptome reference containing all protein coding and long noncoding RNA genes (based on human GENCODE version 32 annotation; downloaded from http://www.gencodegenes.org). The expression counts for each gene (i.e., transcripts per million) in all samples were normalized based on the sequencing depth. Differential expression genes (DEG) were identified using DESeq2 Bioconductor package at a minimum 2 fold change and false discovery rate <0.05 with Benjamini-Hochberg Procedure. The pathway enrichment analysis was performed using a hypergeometric test with Benjamini-Hochberg correction in Ingenuity pathway analysis (Qiagen). False discovery rate <0.05 was used as cutoff to identify significant signaling pathways after treatment.

### Pro-fibrotic biomarker detection with Luminex®

6000 cells/100 μL/well of CCD-18co cells were seeded in 96 well plate. After overnight culture, they were treated with vehicle, 10 ng/mL OSM, 10 ng/mL TNFα, 10 ng/mL IL-1β, 100 ng/mL TL1A, 200 ng/mL IL-36 or 10 ng/mL TGFβ for 16 or 48 hours. After 16 hours, the cell culture supernatant was harvested. After 48 hours, upon collection of cell culture supernatant, the cells were washed once with PBS, and add 35 μL 0.1% tritonx100 in PBS to lyse cells. A customed kit for pro-fibrotic biomarker detection was ordered from R&D systems. The experiment was performed according to the manufacturer’s instruction. Briefly, the supernatant and lysate samples were diluted 1:1 with the diluent provided in the Luminex kit. 50 μL of standard or sample was added to each well. Then 50 μL of diluted microparticle cocktail was added to each well, followed by incubation for 2 hours at room temperature on a shaker at 800 rpm. The plate was then washed by removing the liquid for each well when the plate was set on a magnetic plate separator to allow the magnetic beads to be sequestered, and filling with 100 μL wash buffer, and removing the liquid again. The wash was repeated 3 times. 50 μL of diluted biotin-antibody cocktail was added to each well, followed by covering and incubating for 1 hour at room temperature on a shaker at 800 rpm. Then the plate was washed 3 times. 50 μL of diluted streptavidin-PE was added to each well and incubated for 30 minutes at room temperature on a shaker at 800 rpm. Then the plate was washed 3 times. Then add 100 μL of wash buffer to each well and incubate for 2 minutes at room temperature on a shaker at 80 rpm. The fluorescence signal was read within 90 minutes using a Luminex 200 system.

### Immunofluorescence detection for ACTA2 and COL1A1

1,500 cells/well CCD-18co cells were seeded in CellCarrier-384 Ultra microplates. After overnight culture, cells were treated with a range of concentrations of stimuli, including IL-11, OSM, TNFα, IL-1β, TL1A, IL-36 and TGFβ for 48 hours. Cells were then fixed with 2% paraformaldehyde for 10 minutes. After washing, the cells were permeabilized with 0.25% Triton X-100 in PBS for 10 minutes, following by washing with PBS for 3 times. The cells were blocked with 1% BSA in PBST (0.1% Tween 20 diluted in PBS) for 30 minutes, and then incubated with anti-ACTA2 antibody (Abcam, cat #: ab7817) and anti-COL1A1 antibody (Abcam, cat #: ab34710). Both antibodies were used 1:1000 dilution in 1% BSA in PBST. After overnight incubation at 4 °C, the cells were washed 3 times with PBS, followed by incubation with goat anti-mouse Alexa 594 (ThermoFisher, cat #: A-11032) and goat anti-rabbit Alexa 488 (ThermoFisher, cat #: A-11008) secondary antibodies for 30 minutes. The cells were then washed with PBS 3 times and stained with Hoechst 33342, before being imaged on an Operetta CLS High Content Analysis System.

### Statistics and calculations

Strictly standardized mean difference (SSMD) was used to assess plate quality. It is estimated as

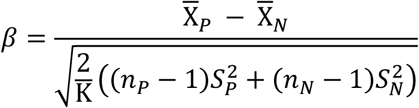

Where 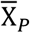 and 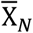 are means, 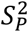 and 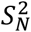 are variances, and *n*_*P*_ and *n*_*N*_ are sample sizes of positive and negative controls, respectively. K ≈ *n*_*P*_ + *n*_*N*_ − 3.48.

For all experiments, statistical analysis was conducted using T-test or one-way ANOVA with Tukey’s post hoc test on Graphpad Prism software with significance of * p<0.05, ** p<0.01, *** p<0.001, **** p<0.0001. Results are presented as means ± SD or means ± SEM, which is specified in the figure descriptions.

## Acknowledgement

We would like to thank our Takeda Development Center of Americas’ colleagues Darren Ruane and Christopher Haines for suggesting the pro-fibrotic stimuli to be used for this study, and Tony Li for CCD-18co cell culture for RNA-seq experiment.

## Author contributions

S.Y., S.R.H. and D.N. concepted and designed the project; S.Y., A.K., M.P., M.M., S.R.H. contributed to the result interpretation and drafted the manuscript; S.Y. performed assay development and high throughput screen experiments; Y.L. participated in high throughput screen experiments; S.Y., Y.L. and Q.W. performed CRISPR/Cas9 experiment; M.M. designed, performed and analyzed scRNA-seq experiment; M.P. performed transcriptomic analysis of CCD-18co datasets and public human colon biopsies datasets; I.I. performed PCA on pro-fibrotic stimuli treated CCD-18co Cell Painting data; J.T. performed PCA on CCD-18co RNA-seq data; J.C. performed Cell Painting primary hit picking with linear predictive model; D.S. and J.H. assisted with laboratory automation and compound management; S.S. and A.E.C. provided input on morphological profiling analysis and its result interpretation, and edited the manuscript.

